# Woodland age, ancient trees, and population size as proxies of genetic diversity

**DOI:** 10.64898/2026.05.16.725641

**Authors:** Efisio Mattana, Nick Atkinson, Irene Martínez-Velasco, Duhyadi Oliva-García, Irene Ramos, Cécile Truchot-Taillefer, Owen Blake, Ted Chapman, Alicia Mastretta-Yanes

## Abstract

Climatic and biogeographic variables are often used as a proxy for tree genetic diversity, but local factors can also influence it. We propose that woodland age, presence of ancient trees, and population size could impact genetic diversity. Using the RBG Kew UK National Tree Seed Project as a study case, we examined how these factors are accounted for during seed collection. We found 42% of tree seed collections come from ancient woodlands and that 8.4% overlap with ancient trees. Sampled forest patches size ranges from few individuals to several thousand. We then carried out a pilot to examine the role of population size on functional traits variation, testing the relationship between population size and seed germination and seedling thermal stress sensitivity in three populations of the *Betula pubescens* Ehrh. complex. We found that seeds and seedlings from larger populations showed higher fitness and stress resistance. Our results highlight the importance of local factors to predict variation in functional traits, relevant for tree resilience. Existing seed collections of native species stored in conservation seed banks offer a valuable resource to explore these factors and improve our understanding of genetic diversity in tree populations, with implications for biodiversity conservation and forestry production.

## Introduction

The establishment of genetically diverse and representative tree populations rely on the identification of provenance zones for the sourcing of reproductive material, often known as *“seed zones”*, “*seed transfer zones*” or “*genetic neighbourhoods”*. Seed zones are often delimited by a combination of environmental factors and expected gene flow distance (Dimon et al., 2026) and are used across the world for conservation and seed production purposes (e.g. Bower et al., 2014; Orquera et al., 2024). For example, the seed zones for native species in the UK (Herbert et al., 1999) are based on major geoclimatic influences and delimited by geological and landform boundaries along the country (Herbert et al., 1999). This official zonation is used for tracking seed collections in commercial production (seed sources), but also to prioritise long term conservation (Kallow, 2024). Although delimiting seed zones by climatic and biogeographic variables accounts for local adaptation to different environments at a coarse scale, it is expected to find genetic differences among populations within a single seed zone (Hamann et al., 2011). However, other than by looking at microclimatic differences, it is difficult to predict how genetic diversity --and the variation in functional traits it confers--will be distributed within seed zones without performing empirical studies, which in turn rely on the availability of seed material.

Here, we argue there are three factors that could positively influence genetic diversity which, however, have been seldom explored by empirical studies: the age of the sampled woodlands, the age of the trees themselves and the size (number of non-clonal mature trees) of the population. To examine how these factors could influence adaptive genetic variation captured during seed collections, we used as study case the UK National Tree Seed Project (UKNTSP) (Chapman et al., 2019) collections stored at the Millennium Seed Bank of the Royal Botanic Gardens Kew (MSB). This is an ideal dataset because it includes hundreds of populations from tens of woody species, with collected seeds maintained separated at maternal level and detailed metadata. We first examined how woodland age, the presence of ancient trees and population size are accounted for during seed collection, by comparing sampling guidelines against what was collected. Then, to exemplify how seed collections can be used to test the role of these factors on functional traits variation, we performed seed germination and seedling stress-tolerance experiments using populations of different sizes of the *Betula pubescens* Ehrh. complex (Downy birch). Since genetic diversity within populations is needed for adaptation and resilience, our findings offer a new perspective for a research agenda supporting seed sourcing strategies in ecological restoration and reforestation programmes.

## Methods and results

### Age of woodlands and trees

Woodland age is expected to influence genetic diversity because old-growth forests which persist in the long-term accumulate mutations and retain local rare alleles, in contrast to recently established populations which may had undergone founder effects and genetic drift (Rajora & Zinck, 2021). In the UK, “recent” or modern secondary woodlands are distinguished from “ancient woodlands”: areas that have been wooded since at least 1600 (1750 in Scotland), and that are the lineal descendants of Britain’s primeval woodland. Although almost always subject to management, their wildlife communities, soils and sometimes old-growth structure have been least modified by humans (Goldberg et al., 2007).

For the above reasons, the UKNTSP collection manual (Kallow, 2024) recommends to “ideally collect from ancient semi-natural woodland”. By examining sampling metadata of the UKNTSP (Figure 1), we found that 499 collections belonging to 55 woody species were collected within ancient woodlands from a total of 1,199 (42%) collections of 92 (60%) species. Therefore, ancient woodlands are not only well represented in the MSB, but the UKNTSP collections provides the opportunity to compare genetic diversity in ancient woodland populations with that of secondary woodlands, and determine whether higher diversity provides functional benefits. This is relevant, because although there are some studies using genetic markers to examine genetic differences between woodlands of different ages (e.g. Rajora & Zinck, 2021), the extent to which they translate into adaptive capacity is still an open question. Beyond research, the availability of materials from ancient woodlands offers the opportunity to support ecological restoration interventions, within and between seed zones, using locally adapted phenotypes or as a source population for climate-smart interventions (Breed et al., 2013).

**Figure 1.**
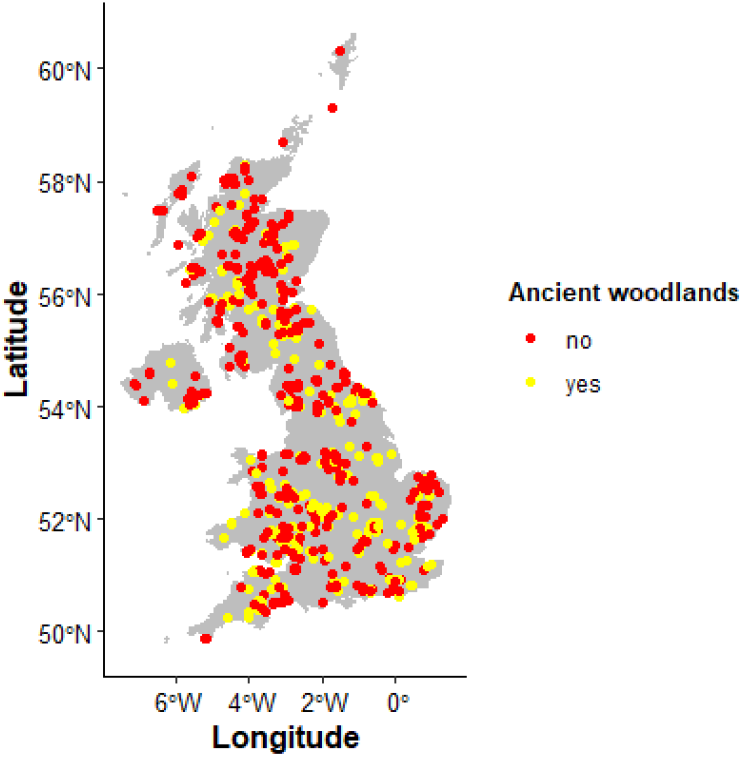
Distribution of MSB tree collections in the UK and overlap with ancient woodlands’ delimitations, based on information recorded in the portfolio data of each collection. UKNTSP data updated to the 15th of May 2025. Collections with no geographic coordinates or identified at genus level only were excluded from the analyses. When coordinates were provided at tree level, they were averaged at population level.

An ancient tree is in the third and final stage of its life. How old an ancient tree is it depends on the species (Nolan et al., 2020). Not all ancient woodlands contain ancient trees, and not all ancient trees are found in ancient woodlands (Nolan et al., 2020). Yet, the presence of ancient trees may influence the present-day population’s diversity because they bring to the mix the diversity of past, lost generations. This is relevant for two main reasons. First, ancient trees have self-evidently demonstrated their ability to withstand large climate variations over time (Cannon et al., 2022). For instance, a thousand-year-old yew has weathered both the Medieval Warm Period and the Little Ice Age. In addition, most secondary woodlands have been intensively planted (often from a narrow gene pool), so it is likely that ancient trees could provide a repository of the genetic variation that once resided in the wider population (e.g. Zhu et al., 2013). Thus, they are living time machines hosting a potential reservoir of genetic diversity, which could support the adaptive capacity of the populations to future extreme climate events. Of course, they could also harbour traits and alleles maladapted to current and future environments. In either case, ancient trees are expected to influence the genetic diversity of the populations they are in, assuming they are still flowering (Cannon et al., 2022; Piovesan et al., 2022).

Ancient trees were not specifically targeted by the UKNTSP project, but we assumed that these collections may contain the genetic pool of an ancient tree if an ancient tree was found in the target population, even if the tree itself was not sampled. For this, we examined the provenance of the UKNTSP accessions against the location of the 18 most common ancient tree species (Downey et al., 2025), according to the ′Ancient Tree Inventory, Woodland trust′ (ATI, https://ati.woodlandtrust.org.uk/) (Figure 2). Although the ATI suffers some biases typical of citizen-science (e.g. volunteers being more active in one region over other, or some species being more favoured), it is an outstanding resource containing hundreds of thousands of verified records. The ATI data confirms that ancient trees often occur outside ancient woodlands, and even outside the species’ natural distribution. For instance, there are several veteran and a couple of ancient Scots pines outside Scotland (Figure 2). Although these trees have been able to survive for hundreds of years outside their native range, these trees were not targeted by the UKTSP (Figure 3) as the project focused on species native ranges (Kallow, 2024).

**Figure 2.**
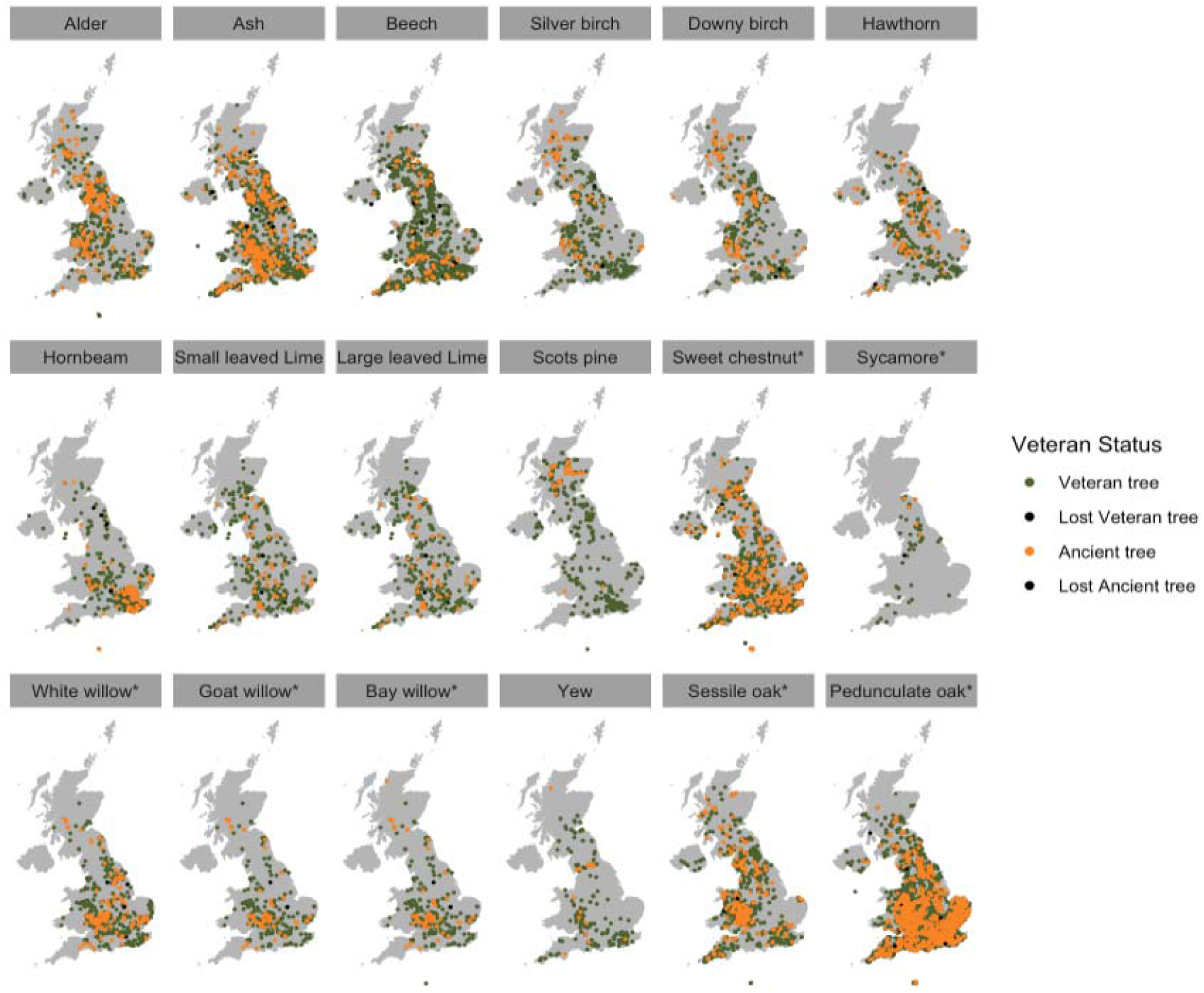
Occurrences of ancient and veteran trees from the most common species reported at the Ancient Tree Inventory (ATI) from the UK. Veteran trees are survivors that have developed some of the features found on ancient trees, they are usually only in their second or mature stage of life, i.e. all ancient trees are also veteran trees (Nolan et al., 2020). Note that the ATI stores only common names, which in birch, lime and willow can be generic for more than one species. Since hybridization is common between species within these genera, we assumed an ancient tree would be contributing to the genetic pool of a population of the same genus. To account for this, in the above maps, a tree labelled with a generic name was duplicated for each species of the genus (e.g. “birch” to both downy birch and silver birch). * = Species with reported desiccation sensitive (recalcitrant) seeds (SER, INSR, RBGK, 2023). ATI data up to the 15th of May 2025.

**Figure 3.**
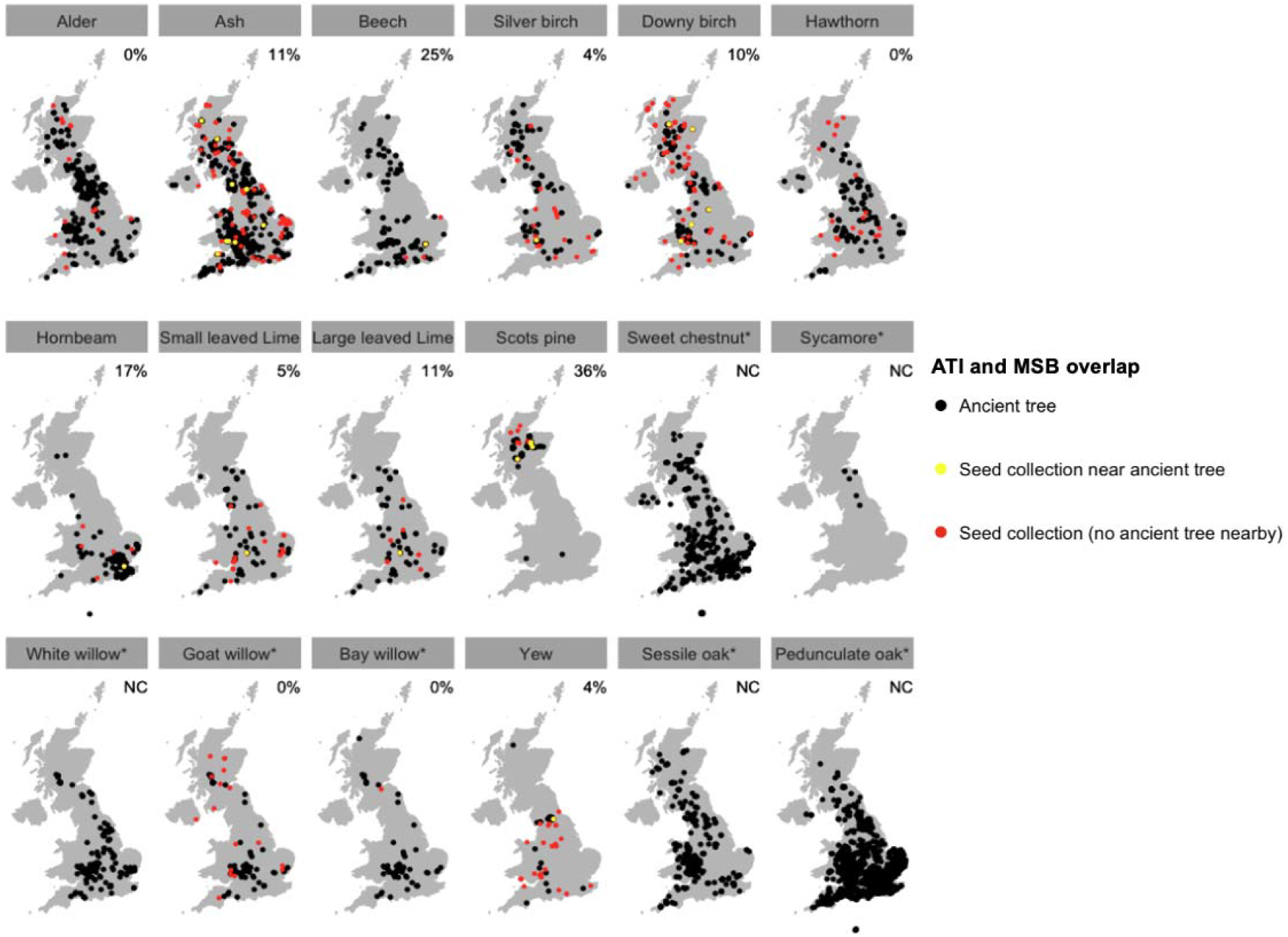
Representativeness of ancient trees in the seed collections from the UK National Tree Seed Project (UKNTSP) stored at the RBG Kew Millenium Seed Bank. We combined two data sources within the MSB: Collections which store geographic coordinates individually for each sampled tree, and accessions which record a single coordinate for several trees from the same locality. To homogenize the data, we averaged the coordinates of individual trees by sampling locality. To examine if seed-collection localities included populations where ancient trees occur, we created a spatial buffer of 2 km radius around each seed-collection locality, and a buffer of 500 m around each ancient tree from the Ancient Tree Inventory (ATI). Common names from the ATI were attributed to the species in the MSB collections according to (Stace, 2019). We considered a positive match (yellow dots) when buffers of the same species intersected. Note that the ATI stores only common names, which in some genera can be generic for more than one species (birch, lime, and willow). In these cases, a single ancient tree could be matched against more than one species (e.g. “birch” to both downy birch and silver birch, or their hybrids). The numbers represent the percentage of seed lots from the MSB that intersect with an ancient tree, NC= Not Collected. * = Species with reported desiccation sensitive (recalcitrant) seeds (SER, INSR, RBGK, 2023).

The data also shows that seven of the most common ancient trees, including oaks (which represent the most abundant and iconic of the UK’s ancient trees), are reported to have recalcitrant (desiccation sensitive) seeds (SER, INSR, RBGK, 2023) and cannot be stored under conventional seed banking conditions (Figure 2). Of the species that can be stored at the MSB, on average 9.5% of their collections came from a population with an ancient tree nearby (Figure 3), totalling 27 seed lots out of the 321 (8.4%) collected by the UKNTSP. The species with the highest percentage of overlap is the Scots pine (36%) and the lowest are alder, silver birch, goat willow and yew (0-4%; Figure 3), which are also poorly represented species on the ATI. Around half (15/27) of the collections overlapping with ancient trees were sampled within an ancient woodland. In summary, the genetic pool of ancient trees is only incidentally represented to a limited extent in the MSB collections.

### Population size and functional traits

The contemporary effective population size (*N*e) is a number that explains the rate at which a population (a group of individuals of the same species mating mostly only among themselves) loses genetic diversity over time by random processes. It is a key concept for conservation biology because populations with a larger *N*e are expected to have higher adaptive potential than populations with a small *N*e, where even reductions of fitness could be expected due to inbreeding (Frankham et al., 2014). In the absence of genetic data, and fulfilling certain assumptions (e.g. no clonality), the contemporary *N*e of a population can be approximated as a fraction of the number of mature individuals using an effective population size / census size (*N*e/*N*c) ratio, most commonly of 10% (Mastretta□Yanes et al., 2024, but see (Allendorf et al., 2024). International biodiversity policies now recognize the importance of *N*e for maintaining populations’ adaptive capacity and countries have started to monitor the size of populations using either genetic data or a *N*e/*N*c-based ratio (Mastretta‐Yanes et al., 2024). Seed collecting manuals, such as the one from the UKNTSP (Kallow, 2024), also acknowledge the importance of population size on maintaining genetic diversity, by setting a target minimum size (4 ha) of forest patches to be sampled. Despite this, little is known on how *N*c and *N*e shape resilience of ecosystem-structuring species such as trees, or how different forms of genetic variation scale up to ecological function.

To address this gap, we set up a pilot study to test whether populations with larger *N*c harbour greater variation in functional traits relevant to coping with environmental change than small populations. We focused on the thermal niche for seed germination and early seedling heat stress sensitivity, both tested under laboratory-controlled conditions. We used seeds from the *Betula pubescens* Ehrh. complex (Downy birch) stored at the MSB, collected from three UK populations differing in population size, including two populations from *Betula pubescens* (“small” and “large” sizes) and one population (“medium” size) of the *B. pendula* x *B. pubescens* hybrid *B*. x *aurata* Borkh. (Figure 4). All populations were collected in ancient woodlands except the smallest one and no ancient trees overlapped with these collections. Seed germination responses to temperature can be modelled by a thermal time approach based on germination rate (GR; (Trudgill et al., 2000). This approach allows the identification of theoretical cardinal temperatures, defining the width of the thermal niche for germination (Figure 4). The thermal niche for germination has been used in several studies to quantify temperature sensitivity during the germination phase of a species/population (e.g., Dayrell et al., 2026). As seeds transition into seedlings, they become particularly vulnerable to high temperatures and water loss (Leck et al., 2008). We expected the thermal window for germination and seedling mass resistance to heath stress to be positively correlated to population *N*c.

**Figure 4.**
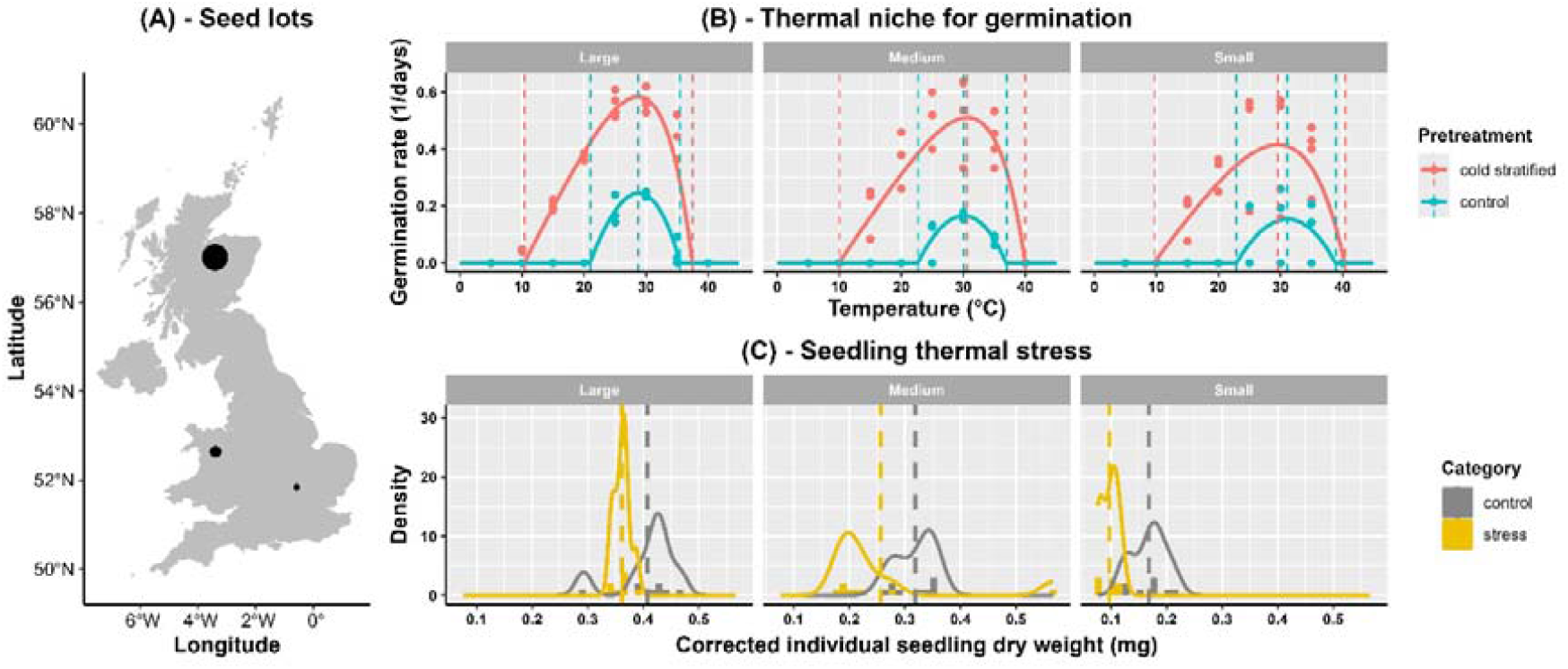
Pilot study carried out on three seed lots of the *Betula pubescens* Ehrh. complex stored at the MSB. (A) Seed lot details. Large population (*B. pubescens*; K:MSB-001010271): ca. 31 km^2^; expected *N*c several thousand trees. Medium population (*B*. x *aurata*; K:MSB-000893613): ca. 2.5 km^2^; expected *N*c > 1000 trees; Small population (*B. pubescens*; K:MSB-000912390): ca. 0.3 km^2^; observed *N*c = 8 trees. Point sizes are proportional to the area of the populations. Population area visually estimated via Google Earth Pro (v. 7.3.6.10441). Further details of each seed lots and natural population are available at: https://herbaria.plants.ox.ac.uk/BOL8/uktreeseed/Home. (B) Thermal window of seed germination. Seeds of four trees from each population were pooled together and incubated in three replicates of 25 seeds each in a gradient of constant temperatures, with and without a cold stratification (one month at 5ºC). The base (Tb), optimal (To), and ceiling temperature (Tc) temperature for germination were modelled following *(Dayrell et al*., *2026)*, estimating the maximum GR and describing the width of the thermal niche for germination (Tc - Tb). (C) Seedling stress responses. 16 cold stratified seeds per population germinated at optimal conditions were incubated at 20ºC. After ten days, half of the seedlings were moved at 45ºC for 3 days to evaluate resilience to heat stress, while the remnant seedlings were kept at the optimal conditions for the same amount of time. At the end of the experiment individual dry seedling weight was assessed. Dry seedling weight was adjusted by subtracting the mean seed weight (0.21, 0.40 and 0.52 mg for the large, medium and small population, respectively), as populations varied in seed mass and seedling mass is known to be highly correlated to seed mass (Simpson et al., 2021), and adding a constant value of 0.5 mg to all seedlings to avoid negative values. Frequency distribution of control and stress seedling was plotted, with dashed vertical line showing mean values of each group.

As previously reported for seeds of this species (Dayrell et al., 2026; De Atrip & O’Reilly, 2007), cold stratification significantly improved seed germination in all populations of *B. pubescens* by breaking physiological seed dormancy, widening the thermal window for seed germination and increasing GR (Figure 4). Contrary to our expectations, we did not detect an increase in thermal window for germination as *N*c increased, with thermal thresholds (Tb and Tc) remaining quite constant among populations in both control and cold stratified seeds (Figure 4). However, in cold stratified seeds, the maximum GR decreased as population size decreased. Slower germination is considered the first symptom of loss in vigour, followed by decreased final germination proportion (Powell, 2022), suggesting that seeds from the large population have a higher fitness. Seedlings (after correcting for seed mass; Figure 4) were larger for larger populations and as *N*c decreased, the difference in mean values between control and stressed plants increased, suggesting a decrease in stress resistance.

These results, suggest that a decrease in ecological functioning may be associated with a reduction in population size, which in turn is expected to decrease genetic diversity. However, the trends described above are the result of a study carried out on a limited number of populations of a single species. Other variables, such as maternal effect (Vivas et al., 2020), environmental conditions (e.g. differences in the average annual temperature or rainfall among populations), and hybridization could have affected seed germination and seedling growth performances.

## Discussion and concluding remarks

Existing and new woodlands must be equipped with a wide range of options to adapt to ever-changing environmental conditions. Maximizing this potential is a significant conservation challenge. Previous efforts have focused on geography and climatic variability; however, woodland age, the presence of ancient trees and population size may also explain patterns of genetic variation within populations. How this translates into adaptive functional traits remains uncertain, but conservation seed banks offer a valuable opportunity to address this question. While these facilities usually aim to capture contemporary genetic diversity (Gargiulo et al., 2025), by storing long-term old collections (Mattana et al., 2025) and, as highlighted here, by sampling in ancient woodlands and around ancient trees, they can also preserve fragments of past genetic diversity. Thus, existing collections provide both metadata and seed materials to test how factors beyond bioclimatic variability influence adaptive traits.

Our findings suggest that population size could have a relevant influence on seed germination and seedling thermal tolerance. Adaptive potential within these traits will be critical in woodlands’ ability to deal with a rapidly changing climate and associated phenological shifts (Dayrell et al., 2026). Further studies should expand this experimental framework across multiple species, populations, and climatic regions to validate these patterns. Incorporating woodland age and the presence of ancient trees as additional variables would help integrate spatial and temporal dimensions shaping genetic diversity. The inclusion of genomic data would be particularly interesting, enabling direct tests of how genetic diversity influences population resilience and how historical and modern factors can be used to predict it. For instance, genomic analyses would allow to calculate the contemporary *N*e and compare it with proxies such as *N*e/*N*c ratio, while also reconstruct historical *N*e (i.e. of past generations), to examine if diversity would be lost given historical and current population trends. Additionally, sampling ancient trees and surrounding populations could reveal whether these trees retain unique alleles otherwise lost, or even to examine the role of somatic mutations within a single ancient tree on generating new diversity.

These lines of research are increasingly relevant as reforestation efforts prioritize high genetic diversity that registered seed sources can provide. For example, Friis et al. (2025) recently found that the distribution of registered seed stands in the UK cover only a small proportion of the bioclimatic variation found in wild populations of ca. 50% of the UK native species studied. To answer these concerns, a network of Genetic Conservation Units (GCUs) is being established in Europe (including the UK), focusing in maintaining high level of genetic diversity of forestry species and providing a source of high-quality seed, protecting the evolutionary potential of forest tree population (EUFORGEN - European Forest genetic Resource Programme - Project”; https://www.euforgen.org/). The factors highlighted here —particularly the roles of ancient woodlands and trees— could be incorporated into such strategy. Once the contribution to contemporary genetic diversity is better understood, their gene pools could help enhance the diversity of reproductive material used in forestry.

We have shown that seed collections stored in conservation seed banks can be used to support new avenues of research. However, our analyses highlighted a key gap: ancient trees are rarely represented in seed collections, unless sampling guidelines explicitly target woodlands that contain them. In addition, considering that many ancient trees occur outside of woodlands, targeting individual trees may be needed too. For example, some of the oldest yews in the UK, are 2,000-5,000 years old, are often found in churchyards and none of them was sampled by the UKNTSP. Citizen science projects like the ATI, and similar initiatives elsewhere (Piovesan et al., 2022) provide detailed distribution data which should be integrated into tree seed collecting planning. Finally, collections made from ancient trees should be explicitly flagged in the collections metadata.

In conclusion, conservation and seed sourcing efforts should look beyond seed zones to represent and maintain genetic diversity. This is true for *ex situ*, but also for *in situ* conservation efforts. To drive evolution, large, long-term persisting populations are essential. Ancient woodlands and trees, which have endured for centuries, already harbour a significant portion of this variation, adding to the reasons for their protection.

## Acknowledgments

We acknowledge the many partners and Royal Botanic Gardens, Kew (RBG Kew) staff involved in the UK National Tree Seed Project (UKNTSP), whose contributions have been essential to this research. We thank the RBG Kew staff who curated the collections and Elena Fouce-Hernandez and Pablo Gomez-Barreiro for their help in setting up the experiments at the RBG Kew Millennium Seed Bank. We thank Roberta Gargiulo, Veronica Reyes-Galindo, Omar Santiago-Clemente and Kew Genetics Group for discussions.

## Conflict of interest disclosure

NA is a director of the Ancient Tree Nursery, a non-profit Community Interest Company focused on genetic conservation of old native trees in the UK.

## Funding

The UK National Tree Seed Project was made possible through the support of players of People’s Postcode Lottery, the Steel Foundation and the John Coates Charitable Trust. The Royal Botanic Gardens, Kew receives grant-in-aid from the Department for Environment, Food and Rural Affairs (DEFRA). Part of the research was conducted during research visits to Kew of IM-V and CT-T, which were particularly supported by Programa de Apoyo a los Estudios de Posgrado (PAEP) from UNAM. Funders had no role in study design, report writing, or article submission.

## Data availability statement

The ′Ancient Tree Inventory, Woodland Trust′ data are available at: https://opendata-woodlandtrust.hub.arcgis.com/datasets/9d2d13b04d654ceb9ba6e0697c1e0c29_0/about. Data with exact coordinates of the UK National Tree Seed Project (UKNTSP) seed collections stored at the MSB is not currently publicly available, however seed lot information with rounded coordinates can be found at: https://herbaria.plants.ox.ac.uk/BOL8/uktreeseed/Home. Seed germination data and scripts from all analyses are available at Dryad repository XXXXX (available upon acceptance).

## Author contributions

EM and AM-Y conceived the idea and designed the research. EM and AM-Y drafted the manuscript, with inputs from NA. EM, IM, DO, IR, CT, and OB collected the data. EM, IM, DO, and CT carried out the experiments. EM, IR, and AM-Y curated and analysed the data. EM, NA, IM-V, DO, IR, CT-T, OB, TC, and AM-Y ′ read, contributed, edited, and approved the final manuscript.

## Notes

https://opendata-woodlandtrust.hub.arcgis.com/datasets/9d2d13b04d654ceb9ba6e0697c1e0c29_0/about

https://herbaria.plants.ox.ac.uk/BOL8/uktreeseed/Home

### Competing Interest Statement

One of the authors of the study, Nick Atkinson is a director of the Ancient Tree Nursery, a non-profit Community Interest Company focused on genetic conservation of old native trees in the UK.

